# Force monitoring reveals single trial dynamics of motor control in a stop signal task

**DOI:** 10.1101/2024.07.05.602195

**Authors:** Surabhi Ramawat, Isabel B Marc, Fabio Di Bello, Giampiero Bardella, Stefano Ferraina, Pierpaolo Pani, Emiliano Brunamonti

## Abstract

The Stop Signal Task (SST) has been the benchmark for studying the behavioral and physiological basis of movement generation and inhibition. In our study, we extended the scope beyond physiological findings related to muscle activity, focusing our analysis on the initial biomechanical state of the effector. By incorporating a force sensitive resistor (FSR), we continuously monitored the force applied by the effector (here the index finger) during a button release version of the SST. This modified task design allowed us to examine both the baseline force before the relevant Go signal was presented and during the covert state of movement preparation. Notably, variations in force over time in response to the Go signal revealed differences across trials where movement was either generated or successfully inhibited, depending on the amount of force during the baseline period. Specifically, higher baseline force was associated with a delayed movement generation, which simultaneously slowed down the force release, facilitating successful inhibition when requested. Our results highlight the influence of biomechanical variables in movement control, which should be accounted for by the models developed for investigating the physiology of this ability.

**NEW & NOTEWORTHY:** Movement involves changing the position of anatomical effectors, like a finger. The initial biomechanical state of an effector impacts movement generation and inhibition. Using the Stop-Signal task, we studied these factors by measuring the force applied to a mouse button before movement onset. Higher initial force delayed movement generation and slowed force release, aiding movement inhibition. This research links behavioral models of action-stopping with movement biomechanics, highlighting the effector’s initial state importance.

## INTRODUCTION

Movement is the process aiming the change of state (position) of anatomical effectors, such as limbs, eyes, or fingers in humans and non-human animals. Mechanisms preserving the position can avoid or facilitate this transition, thereby influencing both movement generation and inhibition. Research has extensively investigated how planned movements could be cancelled, often employing the Stop-Signal task (1, 2). In this task, participants perform a primary action, typically pressing a button with one of their fingers, in response to a “Go” signal, which cues the initiation of the planned movement. Occasionally and unpredictably, a “Stop” signal is presented after the Go signal requiring the participants to inhibit the planned action. Correct and error stop trials are those in which actions are successfully inhibited or executed after the Stop signal presentation, respectively. Analyzing the behavioral performance in this task provides insights into motor control by examining the interaction between action generation and inhibition (2).

Conventionally, studies using the Stop-Signal task have described changes in the dynamic components of the motor responses like those involved in keypresses or dynamometer squeezes to detect movement onset (3–5). In addition, some works have focused on analyzing electromyography (EMG) signals to additionally describe the role of the muscle activations preceding movement initiation and understand movement control more in-depth (6–9). EMG recordings from task-relevant muscles for movement generation reveal subtle changes in activity preceding movement onset (5, 6, 9). Interestingly, bursts of muscle activation have been observed even in correct stop trials when overt movements are not detectable. These activities have been termed partial EMG responses, suggesting a subthreshold activation of the agonist muscle (5, 6). In some cases, the attempts to inhibit the movements were evident in the activation of the antagonist muscles activations (7). Similarly, an effect of inhibition has also been highlighted in the error stop trials observed as a decline in EMG amplitude and prolonged latencies between the peak EMG amplitudes and motor responses (10) suggesting a partial inhibition. Another study has demonstrated an effect of inhibition during the error stop trials through a reduced response force (11).

These findings suggest that the interaction between the processes terminating onto movement generation or inhibition is at play well before the primary behavioral outcome of the Stop Task which is the effector’s action or stillness. Despite these insights, previous studies have largely overlooked the potential influence of the effector’s initial condition, particularly the resting biomechanical state, on movement control. This aspect is well evident in certain movements that perturbate the fully body mass distribution and require anticipatory postural adjustments before execution (12–14). These studies suggest that biomechanical variables linked to the initial posture could influence movement dynamics. However, if these findings also hold true for single effector movements still remains largely unknown. For instance, in a button release task, the amount of force applied on the button (initial muscle tension) at the trial’s outset might determine the latency of responding to a Go signal (15), and possibly the probability of interrupting the response if necessary.

We studied how the initial biomechanical state of the effector can influence motor control by monitoring the force variation on a mouse button over time during a button release version of the Stop-Signal task. By analyzing the signal generated by a force sensitive resistor (FSR) affixed on the button, we estimated the latencies of movement onset and inhibition at a single trial level. Thereupon, we investigated how these variables were influenced by the amount of force applied at the beginning of the trial, as an index of the initial biomechanical state. Finally, we extended these investigations to compare the movement and stopping latencies estimated from the force signal to the ones estimated by muscle activity in a sub-population of the subjects. Our findings bridge the gap between the physiologically plausible models of action stopping and the behavioral models, while introducing the importance of the starting state of the effector(s) prior to the movement preparation.

## MATERIALS AND METHODS

### Participants

A total of 19 right-handed (Edinburgh Handedness Inventory) healthy adult participants (6 females and 13 males) aged between 24 and 48 years (mean: 30 ± 6 years) were enrolled. Experimental acquisitions were performed in accordance with the Helsinki declaration and approved by the local ethic committee.

### Experimental setting

Participants performed a Stop Signal Task (SST) that required a button release response (Figure 1A). They were seated and with their right arm resting on a desk in a comfortable position. Their index finger laid on the left mouse button of a computer where a force sensitive resistor (FSR; Interlink 402) was fixed to detect the force exerted by the finger (Figure 1B). As the figure shows, the FSR was connected to a power supply (V_s_=4.5V) on one end, and a fixed pulldown resistor (R_f_=1 kΩ) on the other. The amplitude of the recorded analog voltage signal corresponded to the applied force as: 1 Volt=2.4 Newtons. The sensor transduced the exerted force into an analog voltage signal proportional to the amount of force on it. The signal was recorded over time at a sampling frequency of 1000 Hz by an acquisition board, and synchronized with the presentation time of the stimuli and behavioral responses (MATLAB; MathWorks, Inc.). In a subgroup of participants (n=6), the EMG activity of the index elevator (extensor indicis) was also recorded by surface electrodes at a sampling rate of 6104 Hz, during the execution of the task.

**Figure 1.**
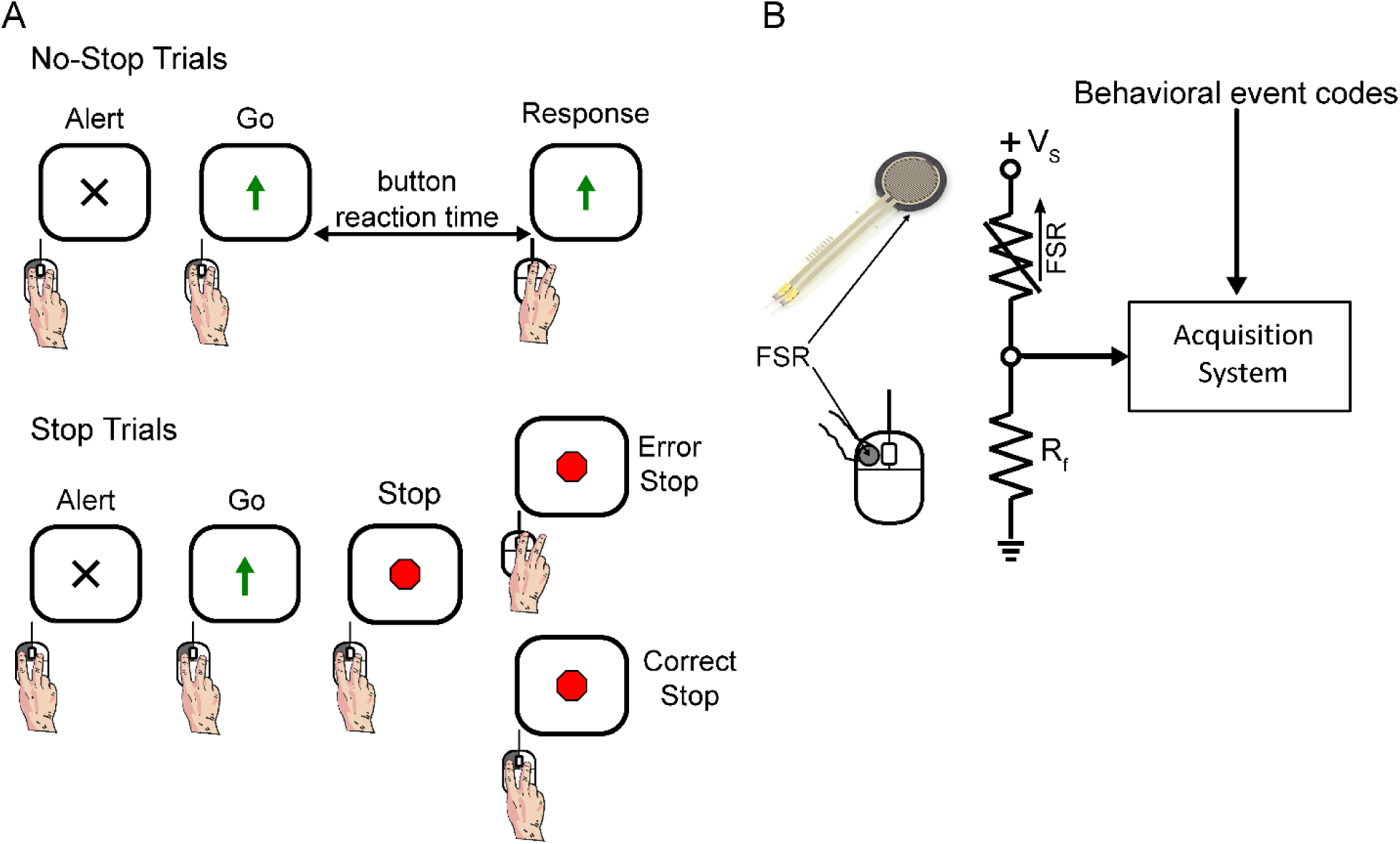
Stop Signal Task and force detection system. A. Schematic of a button release Stop signal Task. Participants were required to lift the index finger in response to the Go signal (Upward arrow) in the No-Stop trials (70% of trials - top panel). Unpredictably, in 30% of trials, the Go signal was followed by a Stop signal (Red Octagon), which indicated to maintain the left button press (Stop trials - lower panel) B. Schematic of the force acquisition sensor circuit and its interaction with the acquisition system. The sensor was powered by a 4.5V voltage source (V_s_) connected in series with the FSR and a 1 kΩ pulldown resistor (R_f_), and the signal was recorded from the point between the two resistors by the acquisition system along with the behavioral event codes sent by the task computer

### Behavioral task

The beginning of each trial was alerted by the onset of a warning signal presented at the center of a computer monitor (Fig. 1A). Participants were instructed to press the left computer mouse button with their right index finger to start the trial. No instruction was provided on the level of force to apply to the mouse button. Following a variable time between 1000 and 1500 milliseconds, a Go signal (an upper pointing arrow) replaced the warning signal, requiring the participants to lift their index finger as fast as possible. This response was expected in 70% of the experimental trials (No-Stop trials). In the remaining 30% of the trials, a Stop signal (a red octagon) replaced the Go signal after a variable delay (the Stop Signal Delay or SSD), requiring participants to keep the mouse button pressed. The trials in which this request was accomplished were named Correct Stop trials, while the Stop trials in which the button was released were labeled as Error Stop trials. A staircase procedure adjusted the length of the SSD by adding 50 ms to the latest SSD if a Correct Stop trial was performed, and decreasing the SSD by the same value after an Error Stop trial. The initial SSD of each recording session was set at 50 ms. The SSDs computed by the staircase procedure was randomly intermingled with two fixed SSDs, 100 and 400 ms in 30% of the total Stop trials, respectively. The fixed SSDs were introduced to obtain a higher number of Stop trials in the first and last quartile of the overall SSD distribution to explore the impact of the length of the SSDs on sensor-based measures of the reaction time to the stop signal. An upper time limit for the button release was set at 1000 ms to discourage waiting for the Stop signal. Trials in which the response was generated before the Go signal presentation or the button release was not initiated until the upper time limit was achieved were aborted with auditory error feedback and excluded from the analysis.

Every participant completed a total of 1200 trial divided in 3 blocks of 400 trials each within the same session. The rest period between consecutive blocks depended on when the participant felt ready to start the next one. The initial SSD of each block after the first one, was based on the final SSD value from the block before it.

### Identification of partial Error Stop Trials

Previous studies recording EMG activity in SST identified a proportion of Correct Stop trials with an initial increase of antagonist muscle activity after the presentation of the Go signal, followed by a decrease after the Stop signal presentation (5, 9, 16). In these trials, named partial Error Stop trials, although no overt response was detected, the activation in response to the Go signal was interpreted as the evidence of a muscle preparation for the following movement partially influenced by the Stop signal presentation. Similarly, in the present study, we observed Stop trials that showed a change in force detected by the sensor after the Go signal that was not followed by a movement of the mouse button (Fig. 2). To identify these trials, we computed in each Correct Stop trial the velocity of the FSR signal over time. If, during the trial, the negative magnitude of velocity was maintained above the average value computed in the 200 ms preceding the Go signal minus two standard deviations, the trial was labeled as Correct Stop. On the contrary, if the velocity crossed this threshold and then turned back above this limit, the trial was identified as partial Error Stop (Fig. 2). In searching for these trials, we evaluated whether different amounts of partial error trials could be linked to longer or shorter SSD. Since we did not detect differences between the two fixed SSDs, we did not further explore this hypothesis.

**Figure 2.**
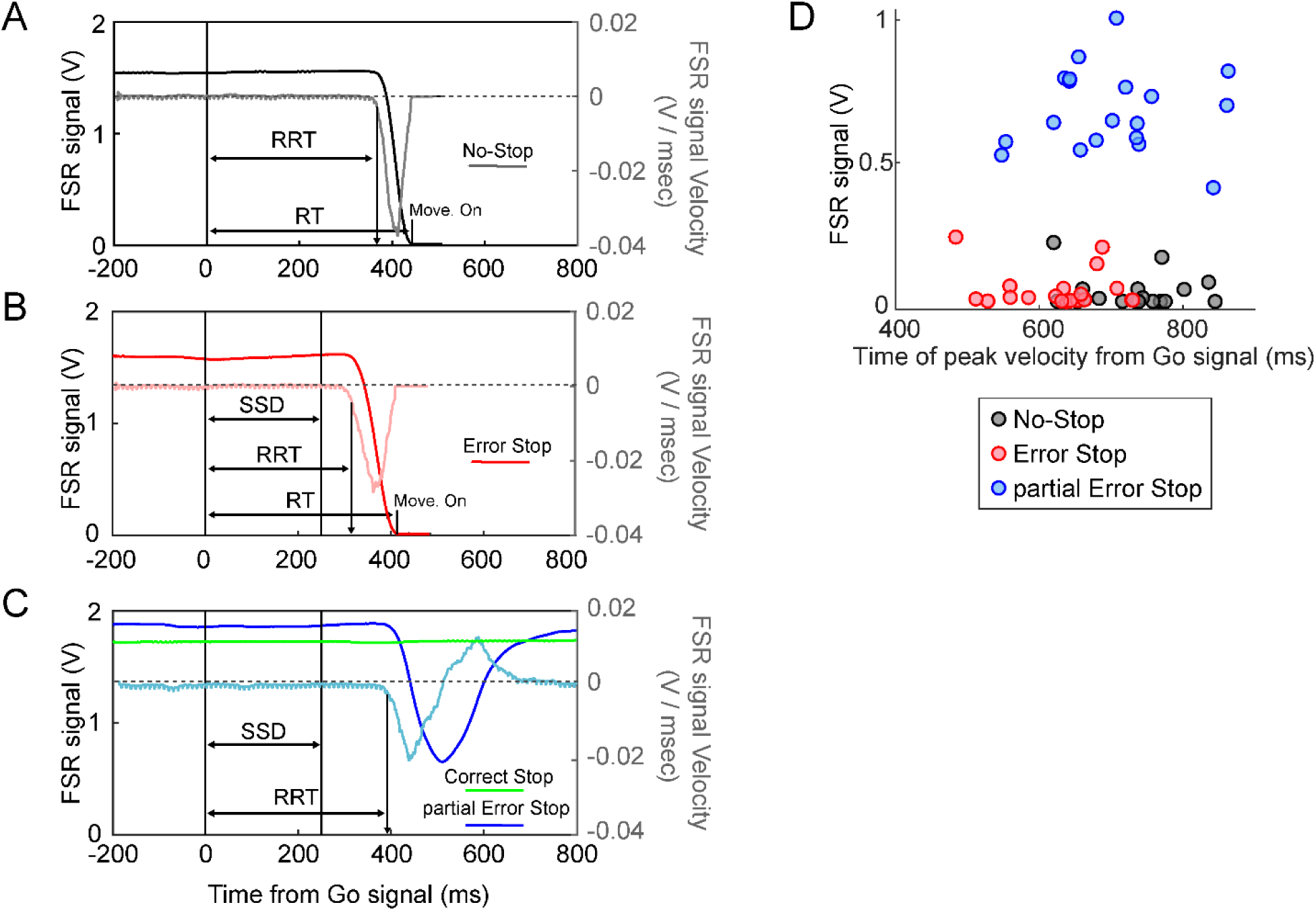
Force sensor signal dynamics across trial types. Task execution is characterized by different relationships between finger force release and the motion detected by releasing the mouse button. The figure shows the temporal dynamics of the voltage equivalent of the force detected by the FSR during single example trials after the Go signal, and the instantaneous velocity of this analog signal (in lighter-colored curves with magnitudes indicated on the right y-axis) for the different trial types: A. No-Stop trials, B. Error Stop trials, and, C. Correct and partial Error Stop trials. The Release Reaction time (RRTs) across trials is detected as the onset of the release of the force after the Go signal, while the RTs are detected as the movement onset (button release) from the Go signal. D. Value of the detected force at the time of the negative peak of the velocity curve for each subject during the No-Stop, Error Stop, and partial Error Stop trials. A higher magnitude of the FSR signal during partial Error Stop trials signifies a slower decline of the applied force, which was reinstated resulting in no overt response.

### Estimation of parameters of movement initiation and cancellation

The Stop Signal Reaction time (SSRT) is a critical measure to probe into the interaction between the Go and the Stop processes during an SST. Conventionally, the SSRT is calculated by fitting the horse race model to the observed behavioral data (1). In this study, we used the velocity of the force release to estimate several parameters of the movement initiation and cancellation as the release reaction time (RRT) to the Go signal, and sensor-estimated SSRT in the partial Error Stop trials.

In No-Stop trials, Error Stop, and partial Error Stop trials, we estimated the RRT as the time when the negative velocity corresponding to the force release fell below three standard deviations of the average velocity in the 200 ms preceding the Go signal onset. These RRTs were then compared to the reaction time (RT) in No-Stop and Error Stop trials observed as the latency of finger movement signaled by the mouse button release. (Fig. 2, panels A and B).

In the partial Error Stop trials, we calculated the sensor-estimated SSRT as the latency between the Stop signal presentation and the time when the instantaneous velocity of the force signal reached its negative peak (as illustrated in Fig. 6A). To assess the reliability of the sensor-estimated SSRT, we compared this measure to the one calculated from the horse race modelling approach. In doing so, we applied the integration method (1) using two procedures; in the first case, by considering the average SSD and the average probability of response; and in the second case by considering the three central SSDs for each subject, that provided a probability of response closer to 50%. The fixed SSDs were also included in the SSRT estimation. We then computed the SSRT for each of the three SSDs and averaged the values. Further exploring the effect of the initial state of the effector on the motor initiation and inhibition, we compared the RRTs and the SSRTs obtained from No-stop and partial Error Stop trials from the first and last quartiles of baseline force distribution within these trial types across subjects. We also assessed the probability of performing a correct stop trial ‘p(stop)’ in the first and the last quartiles of overall baseline force distribution within each session.

To compare the rates of FSR signal decay across different trial types, we calculated the average velocity of the signal during the epoch ranging from the onset of the force release to its completion in No-Stop, Error Stop, and partial Error Stop trials. We checked if the different lengths of the trials affect this average velocity by additionally calculating the average velocity from the initial 50 ms after the onset of force release.

### Evaluation of the effect of force on movement control

In a second experiment, we tested if the magnitude of force on the mouse button influenced the parameters of movement initiation and cancellation by asking 10 participants of our sample to perform further blocks of trials in two different conditions. For these blocks, we defined a lower and an upper force threshold based on the range of values of the first and last quartile of distribution of the baseline force recorded in the first experiment. In the first high initial force condition the subjects were asked to start each trial by pushing the button with the index finger with a FSR signal magnitude of at least the upper force threshold; while in the other low initial force condition, they were required to start the trials with the signal level less than the lower force threshold. Acoustic feedback signaled whether the exerted force was below or above the defined limit respectively, helping them to maintain the requested level of force. By analyzing the time course of the force signal on each trial (see previous methods), we quantified the performance metrics like the number of trials, RRTs and SSRTs for each force condition.

### Processing of muscle activity signals for estimating the parameters of movement initiation and cancellation

The muscle signal was rectified, down-sampled at 1000 Hz, and smoothed by a moving window of 30 ms. To detect the muscle reaction time (muscle RT) to the Go signal in No-stop and Error stop trials, and the SSRT to the Stop signal in partial Error stop trials, we applied an algorithm similar to the one used for the force signal in respective trial types. Specifically, we computed the velocity of the agonist muscle contraction and estimated the muscle RT as the latency between the Go signal and the time at which the velocity crossed its average value computed in the 200 ms preceding the Go signal plus 2.5 standard deviations (mean Baseline Velocity + 2.5×STD). The muscle SSRT was estimated in the partial Error Stop trials as the latency between the Stop signal onset and the time of the peak velocity of the activity.

### Statistical Analysis

The analysis was performed using customized scripts in MATLAB (The MathWorks, Inc, version 2023a). To test for the across-group differences, One-way ANOVA and paired t-tests were employed. However, to see the effect of the SSDs and the Stop trials types on the level of force we used a two-way ANOVA with interaction. Also, for the control experiment, all the comparisons for the within task and across task conditions (low initial force vs high initial force) were performed using a two-way ANOVA. We performed the regression analysis by fitting the data to a linear equation. We obtained the coefficients for the correlation analysis using Pearson’s correlation (r). The post-hoc comparisons after obtaining a significance were performed by Newman-Keul’s method. All the results are reported as Mean±Standard Deviation. For our results, a p-value p<0.05 was considered significant.

## RESULTS

### Specific dynamics of force release characterizes movement onset

In No-Stop trials, as shown in Fig. 2A, a sharp decrease in force was observed until a total release before movement generation. This decrease is characterized by a bell-shaped velocity curve. We estimated the release reaction times (RRTs; see methods), and we found that it occurred always before the behavioral reaction times (RTs) calculated from the movement onset detected by the mouse button (451±64 ms and 572±66 ms; t(18)= −21.1, p<0.001). This relationship was confirmed also in the Error Stop trials (Fig. 2B), where a similar delay between the RRTs and mouse button release RTs was observed (359±70 ms and 479±71 ms; t(18)=-23.5, p<0.001). In the partial Error Stop trials, a decrease in force was initiated, which was then interrupted and reversed its course before causing the mouse button displacement, allowing us to measure the RRT in response to the Go signal; while in the Correct stop trials, force release and mouse button lift were not detected (Fig. 2C). Using the velocity-based threshold (see methods for details), we quantified 150±22 Error Stop, 42±15 partial Error Stop and 129±21 Correct Stop trials across subjects.

Thus, the data shows that movement generation is characterized by a rapid release of force followed by the movement of the mouse button, although in some of the trials, the release of force could be resisted and as a result, no movement was detected by the mouse button. These observations suggest that there exists a threshold in the force dynamics, which, if crossed, will lead to a displacement of the mouse button. Investigation of this phenomenon would allow us to identify a point of no return for motion generation in this experimental context, that is, a point in the force release dynamics beyond which the motion of the mouse button is necessarily generated.

To this end, we compared the different types of trials in which a deflection in the applied force was detected after the Go signal (No-Stop, Error Stop and partial Error Stop trial). We observed a significant difference in the force values recorded at the moment of peak velocities between the trial types (Fig. 2D; One-way Anova: F(2,36)=230.18; p<0.001). A Newman-Keuls post-hoc comparison revealed that the mean FSR signal values at peak velocities in the trials characterized by movement generation (No-Stop: 0.044±0.057 V and Error Stop: 0.055±0.066 V) and the trials resulting in successful movement inhibition (partial Error Stop: 0.68±0.14 V) were significantly different. In the No-Stop and Error Stop trials, the FSR signal was detected to be less than 0.3 V at the moment of minimum peak of velocity, however, in partial Error Stop trials, the signal level was maintained at a level higher than 0.4 V (Fig. 2D).

In general, we found that movement generation is anticipated by a release of force that triggers the movements detected by the mouse once the force drops below a specific level.

### The force level before the Go signal varies according to the type of trial

Once we established that force release is a key component of movement generation, we asked whether there is a relationship between the level of FSR signal before the Go signal and performance (i.e., movement generation or inhibition). To this aim, we compared among the different types of trials the level of force on the button in the time before the Go signal (baseline force). We found that the baseline FSR signal level was higher on average in the partial Error Stop trials and the Correct Stop trials than in the Error Stop and No-Stop trials (Fig. 3A; One-way Anova: F(3,72)= 10.03; p<0.001). Post-hoc comparisons detected a significant difference between the baseline FSR signal detected in No-Stop trials in comparison to Error, Correct, and partial Error Stop trials (p<0.05). Moreover, the baseline FSR signal in partial Error Stop trials was also significantly higher than the Error Stop trials (p<0.05). Thus, the level of baseline force seems to participate in determining whether or not a movement will be inhibited.

**Figure 3.**
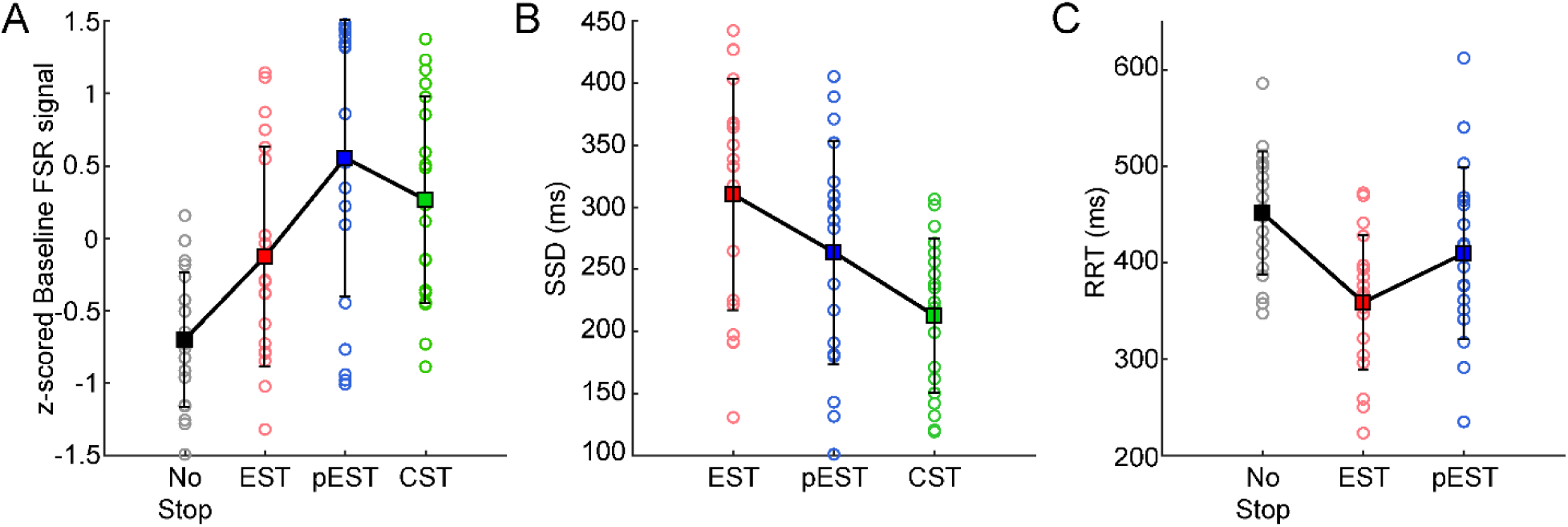
Performance metrics obtained by FSR in different trial types. A. The normalized mean FSR signal detected by the sensor during the baseline period (200 ms before the Go signal) for No Stop (gray circles), Error Stop (EST, red circles), partial Error Stop (pEST, blue circles), and Correct Stop trials (CST, green circles) for each subject. B. The mean Stop Signal Delay (SSD) across the different Stop trial types, i.e., Error Stop, partial Error Stop, and Correct Stop trials for each subject. C. Release reaction time detected through the decline in the force sensor signal as a response to the Go signal in No-Stop, Error Stop and partial Error Stop trials for each subject. The squares and the vertical bars indicate the mean and the STD across the subjects, respectively.

According to the horse race model, an increase in the SSD corresponds to a higher probability of responding in Stop trials, and the longer the RTs in the No-Stop trials, the lower the probability of responding in a Stop trial. Consistent with this model, we found that Error Stop trials were more likely to occur when the Stop signal was presented at longer SSDs (310±93 ms), whereas the Correct Stop trials were observed at shorter SSDs (213±62 ms). The SSDs at which partial errors were detected were those with values between these two extremes (264±90 ms; Fig. 3B; One-way Anova: F(2,54)= 3.168; p<0.01). Furthermore, we observed higher RRTs during the No-Stop (451±64 ms) trials compared to Error Stop trials (359±70ms), while the RRTs obtained in partial Error Stop (409 ±89 ms) trials were on average lower than No-Stop RRTs, but higher than Error Stop RRTs (Fig. 3C, One-way Anova: F(2,54)= 7.188; p<0.01). Newman-Keuls post-hoc comparisons revealed significant differences in the SSDs and RRTs across different trial types (p<0.05).

These findings suggest that both, the level of baseline force amplitudes and the SSDs affect the performance in the stop trials. To study in more detail the specific contributions of both of these parameters on movement control, we evaluated whether the occurrence of the partial Error Stop trials was primarily related to the shorter SSD durations or the baseline force levels. We ran a two-way Anova (factors: SSD and Stop trial type) and we detected a significantly higher level of FSR signal (F(1,9)= 4.64; p=0.02) in partial Error Stop trials than in Error Stop trials, as shown in Fig. 4. Significant effects of SSD (F(9,288)= 4.64; p=0.39) or interaction (F(9,288)= 0.29; p=0.98) on the baseline FSR signal were not detected. This result suggests that the force exerted on the button at the beginning of the trial might have an influence on the outcome of motion inhibition per se, independent of the SSD.

**Figure 4.**
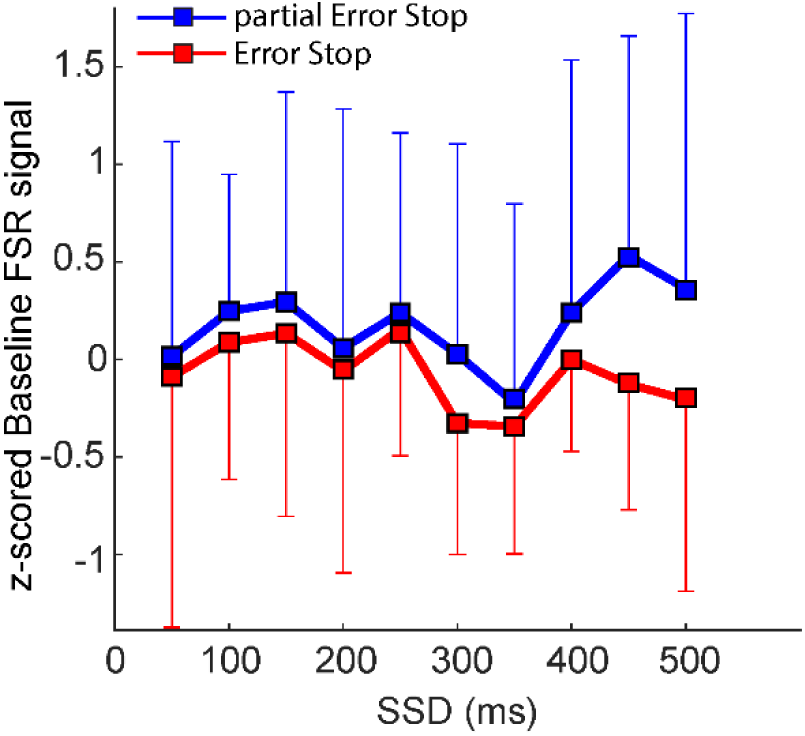
Relation between the baseline force and the SSDs in the Stop trials. The figure shows the mean baseline FSR signal at each SSD characterizing the partial Error Stop (blue plot) and the Error Stop (red plot) trials. The plots show that the baseline force was significantly higher in the partial Error Stop trials across all the SSDs, specifically for the longer SSDs. The squares and the vertical bars represent the mean and the STD across subjects.

Correct and partial Error Stop trials were the ones with higher degrees of force on the mouse button and occurred for shorter SSDs. On the contrary, a lower degree of force and longer SSD were associated with Error Stop trials. Interestingly, in comparing Error Stop and partial Error Stop trials we observed higher baseline force, shorter SSDs, and longer RRT in partial Error than the Error stop trials. To conclude: the higher the pre-applied force, the higher the probability of having an inhibited, partial Error Stop trial following the Stop signal.

### The response times in No-Stop trials are affected by the level of force before the Go signal

Once we established that the level of force affected the performance in stop trials, we explored whether a relationship between the magnitude of baseline force and the RRTs could be detected. To answer this question, we identified the first and last quartile of the baseline FSR signal level distribution across all the No-Stop trials (200ms before the Go signal) overall participants, and then compared the RRTs associated with these portions of the distributions. The average FSR signal magnitude was approximately 0.7and 1.3in the first and fourth quartile, respectively. As detailed in Table 1, the average RRTs were significantly longer when the initial force applied on the mouse button was higher, compared to when the button press was initiated with a lower force (510 Vs 476; p<0.01).

**Table 1.** Average RRTs, SSRTs and the probability of correct stop trials ‘p(Stop)’ calculated within the first and the last quartiles of baseline force distribution across same and different trial types.

|  | First Quartile | Last Quartile | Statistics |
| --- | --- | --- | --- |
| **No-Stop Trials** |  |  |  |
| **Baseline FSR Signal** | 0.69±0.25 V | 1.32±0.36 V |  |
| **RRT** | 476.03±244 ms | 510±252.23 ms | $t(1554)=1.96$, $p=0.006$ |
| **Partial Error Stop Trials** |  |  |  |
| **Baseline FSR Signal** | 0.69±0.25 V | 1.36±0.37 V |  |
| **SSRT** | 172±34 ms | 177±26 ms | $t(196)=-0.18$, $p=0.85$ |
| **All Trials** |  |  |  |
| **Baseline FSR Signal** | 0.65± 0.2 V | 1.34± 0.3 V |  |
| **p(Stop)** | 0.50±0.07 | 0.55±0.07 | $t(18)=-2.7$; $p=0.012$ |
| **pEST Trial Count** | 8±4 | 12±6 | $t(18)=-2.8$, $p=0.009$ |

These data confirm that the force applied on the button before the Go signal influences the response times related to the movement execution during the No-Stop trials.

### Trial-based estimate of SSRT by the dynamic of force release in partial Stop Errors

Extending these findings to the partial Error Stop trials is particularly interesting because these trials are characterized by a pattern of FSR signal that indicates an initial release of force followed by an increase in force. The restoration of the applied force prevented the generation of overt movement detectable by the mouse button. This dynamic is possibly a reflection of the inhibition process at play to prevent movement generation. As in EMG studies (16), here we obtained an estimate of the SSRT by capturing in partial Error Stop trials the time at which the Stop signal elicited a correction on the ongoing movement as depicted in Figure 5A (refer to methods for details). We obtained a distribution of the sensor SSRT estimates from each participant as represented in the Fig. 5B (one example subject). Consequently, we computed the SSRT using two approaches that relied on the horse race model, as described in the methods. Both of these approaches provided comparable estimates of the SSRT (SSRT Method 1 = 262±63; SSRT Method 2 = 259±62; t-test: t(18)= 2.10, p=0.76). Since the two behavioral estimates of SSRT were not significantly different, we averaged them to obtain a single behavioral estimate of the modeled SSRT (SSRTm). We then compared SSRTm with the sensor estimated SSRT (Fig. 5C). The figure shows that the SSRTm was significantly longer (260±60 ms) than the sensor-estimated SSRT obtained from the FSR signal dynamics (172±24 ms); (t-test: t(18)= 2.10, p<0.001). However, the SSRTm and the sensor-estimated SSRTs were highly and significantly correlated (Pearson correlation coefficient r= 0.71; p<0.001), instating the latter to be a reliable measure of the inhibitory control.

**Figure 5.**
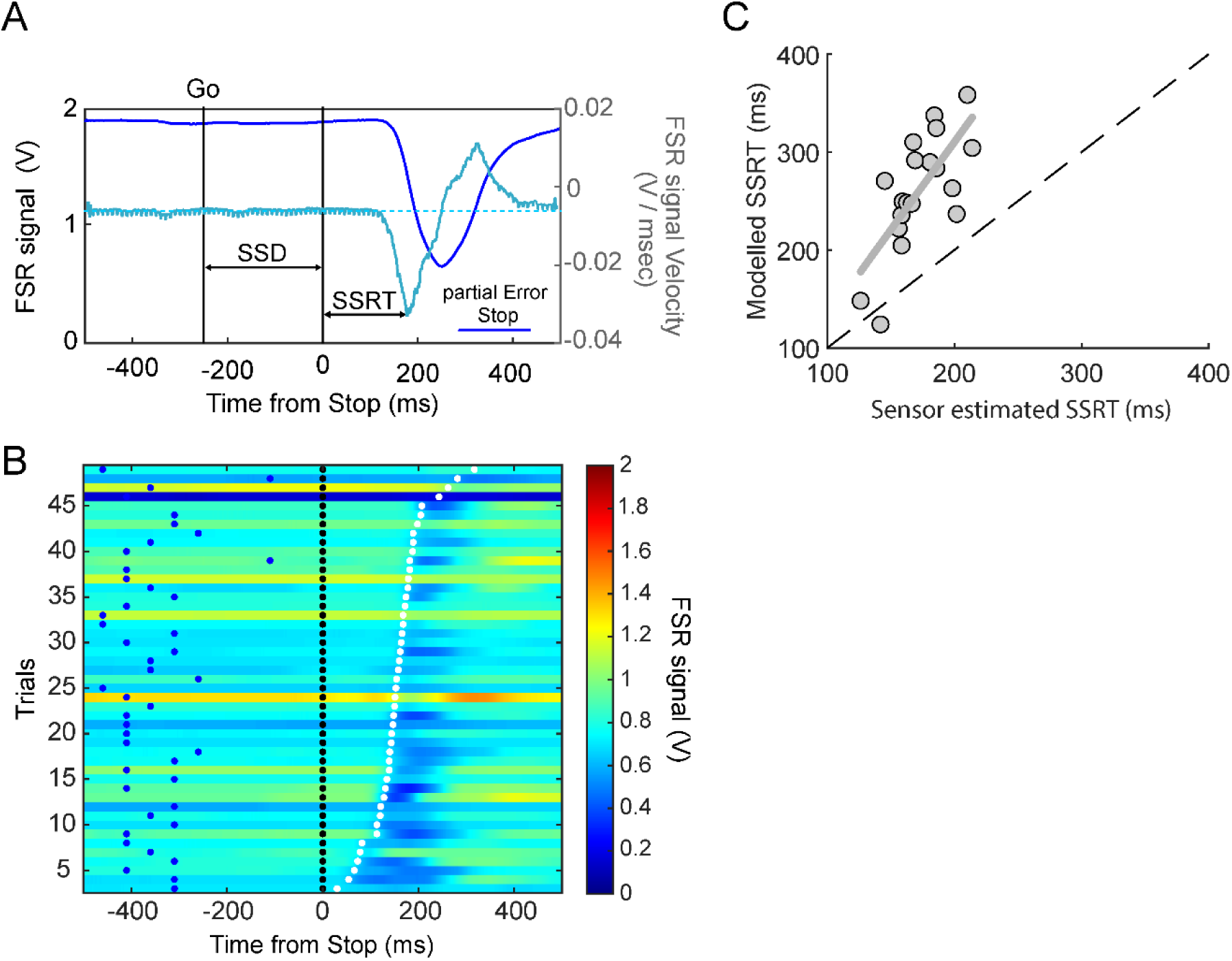
Force release dynamics in movement inhibition. A. The plots show the time course of the analog FSR signal (dark blue) and the corresponding instantaneous velocity (light blue) during an example Stop trial detected as a partial Error Stop trial by the FSR signal. The Stop Signal Reaction time (SSRT) is estimated as the time of crossing the threshold in the velocity profile resulting from the deflection and the restoration of the FSR signal. B. The color map shows the FSR signal amplitude detected for all the partial Error Stop trials during an example session from a single subject. The trials are represented along the y-axis, and the time from the Stop signal (black dots) is represented on the x-axis. The blue and the white dots indicate the Go signal and the sensor estimated SSRT, respectively, for each trial (as shown in panel A). C. The figure represents the SSRTs estimated using the force sensor (x-axis) versus the SSRTs estimated using the model for each subject. The two estimates are strongly correlated, while, the sensor-estimated SSRTs are shown to be faster than the model estimates SSRTs.

Following the analysis of RRTs and trial type, we asked whether the force before the Go signal also affected the length of the SSRT. To this end, we calculated the values of FSR signal corresponding to the first (0.69 V) and the last (1.3 V) quartiles of the baseline force in the partial Error Stop trials for each subject. Differently from the RRTs, we found that, at least in partial Error Stop trials, the level of force before the Go signal did not affect the duration of the sensor estimated SSRTs overall subjects (172 ms Vs 177 ms; See Table 1 for details). However, when looking at the distribution of baseline force across trials in each session, we found that the probability of inhibiting to the stop signal, or to perform a correct stop trial was significantly higher for the higher baseline force (p(stop)= 0.55) in comparison to the lower baseline force (p(stop)= 0.50; p<0.05) across all the subjects. Additionally, the average number of detected partial Error Stop Trials was also higher for higher baseline force in comparison to the lower baseline force (Table 1).

Furthermore, we evaluated whether a preliminary sign of successful inhibition, different from the dropping below the FSR signal threshold, could be found in partial Error Stop compared to Error Stop trials.

Indeed, both trial types were characterized by an initial release of force that, in partial Error trials, was followed by an increase that prevented the lifting of the finger. We found that force dynamics was also characterized by a specific pattern of force release velocity: indeed partial Error Stop trials compared to No-stop and Error stop trials were characterized by a lower magnitude of average release velocity (Fig. 6A; F(2,54)= 25.22; p<0.001; post-hoc p<0.05). Additionally, we calculated the average velocities in the fixed epochs of initial 50 ms, which also resulted in the same pattern (No-stop: 0.0048±0.0016 V/ms, Error stop: −0.00466±0.0016 V/ms and partial Error Stop: −0.0024±0.0008 V/ms). Even this parameter was significantly higher for partial Error Stop trials than the No-stop and Error Stop trials (F(2,54)=16.51; p<0.001). This shows that in the partial Error Stop trials, a stopping process is possibly at play well before the observed release of the force on the mouse button. Moreover, the average baseline FSR signal highly correlates with the average velocity of FSR signal decay across participants (Fig. 6B) for all three trial types (Pearson’s Correlation Coefficient-No-Stop trials: r=-0.88; p<0.001, partial Error Stop trials: r=-0.99; p<0.001, Error Stop trials: r=-0.89; p<0.001). A post-hoc comparison revealed significant differences between the coefficient of linear regression representing the slopes of the lines corresponding to partial Error Stop trials (−0.0028), in comparison to the No-Stop (−0.0052) and Error Stop trials (−0.0053; p<0.01). These observations emphasize the effect of the biomechanical state before the Go signal and its immediate variation in response to the resulting movement or inhibition.

**Figure 6.**
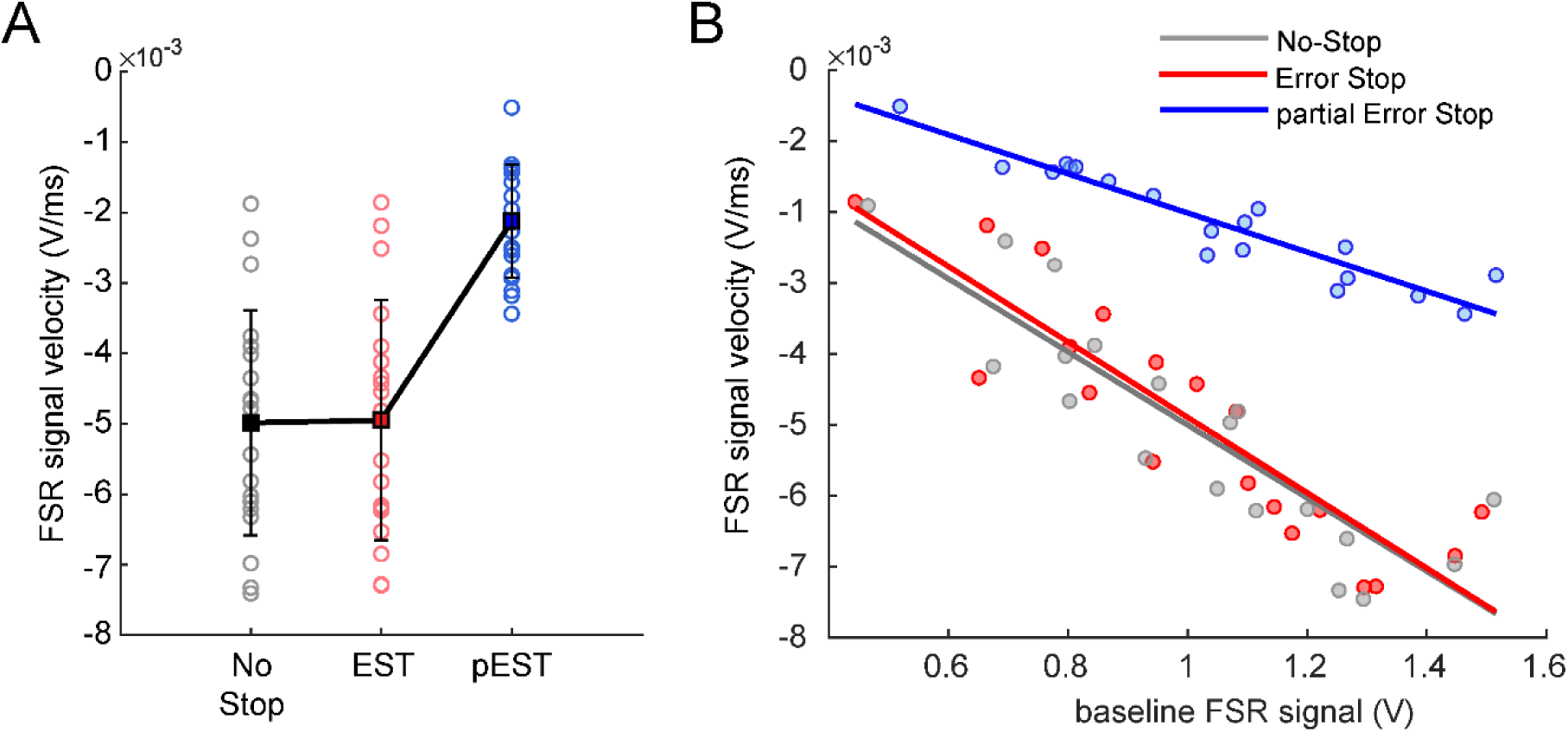
Different FSR signal-velocity profiles across different behavioral outcomes. When the force is released (reflected by the decay in FSR signal) as a result of the Go signal presentation, the average velocity of this signal over time, represented as negative values along the y-axis, reflects the motor outcome. A. The higher negative magnitude of the velocity corresponds to a faster force release resulting in a movement generation, as shown in the plots for each subject for No Stop (gray) and Error Stop (red) trials, while a lower negative magnitude corresponds to a slower force release, which did not result in a movement generation in partial Error Stop trials (blue). B. The plot represents strong correlations (p<0.001) between the baseline FSR signal values and the average velocities of force release in different trial types for each subject.

### Prescribed level of force before the Go Signal affects movement generation but does not affect movement inhibition: A Control Experiment

To explore more in-depth the effect of the force applied before the Go signal on the performance, we ran a second experiment. In this experiment we asked 10 participants from the original sample to perform the task by starting the trial with either a high (> 1.3 V) or a low (< 0.6 V) prescribed force (FSR signal magnitude) on the button. Figure 7A illustrates the distribution of FSR signal magnitudes as a measure of exerted forces for the two task conditions across the participants for the different trials of the task. We confirmed that both, the force level and the trial type significantly affected the RRTs (Fig. 7B; Two-way Anova: main effect of force condition: F(1,9)=4.74; p=0.057) and the type of trial (main effect of trial type: F(2,18)=11.67; p<0.001). Even though the effect of force levels on RRTs was at a first test marginally significant due to the huge variability across the subjects, we found a highly significant modulation of force level and trial types on the normalized scores (main effect of force condition: F(1,9)=11.00; p<0.01) and main effect of trial type: F(2,18)=22.04; p<0.001).

**Figure 7.**
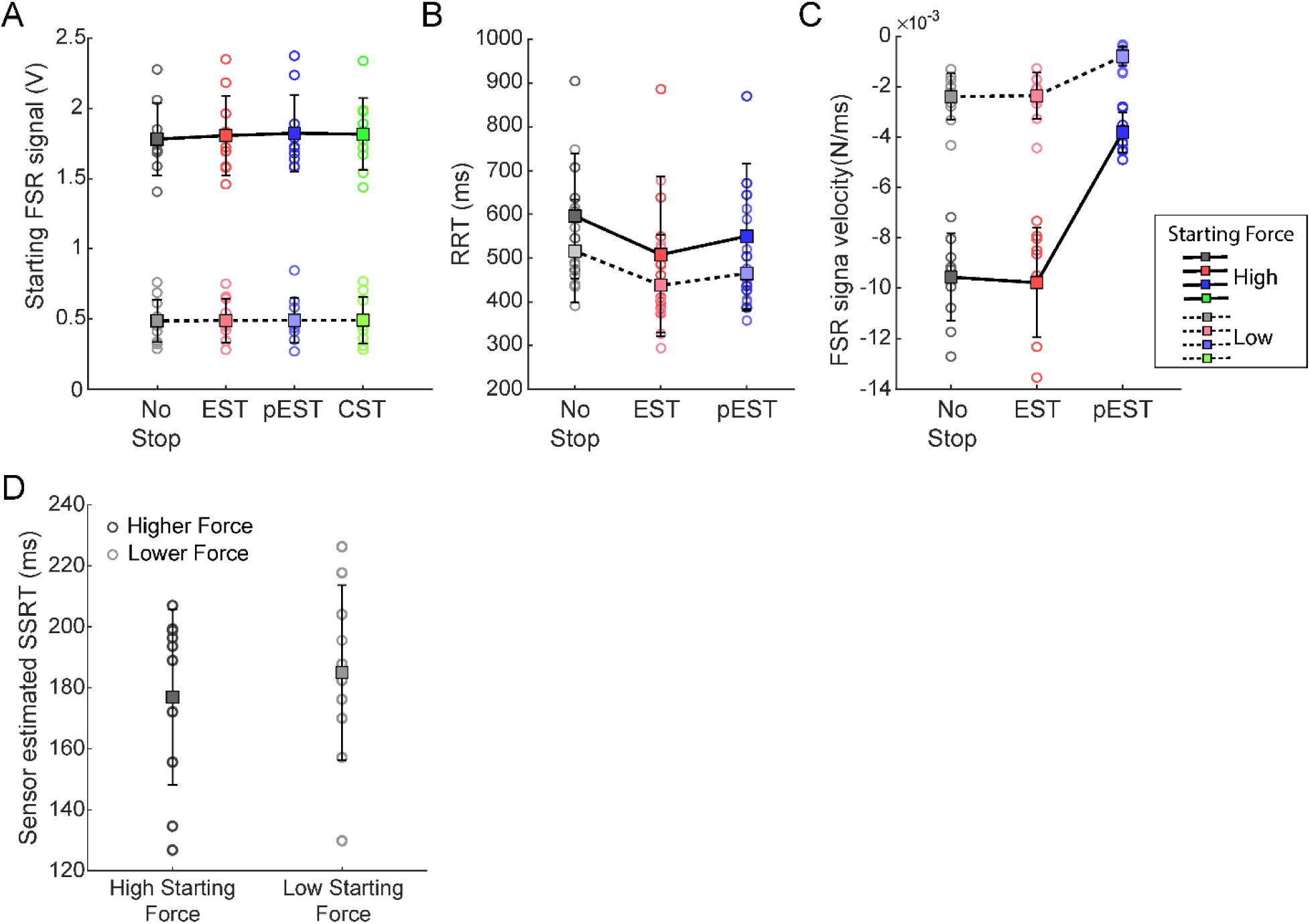
Effect of starting force on movement initiation and inhibition. A. The plot shows the distribution of starting force across a subsample of 10 subjects during the two conditions, low starting force (FSR signal<0.6 V; lighter colors and dashed line) and high starting force (FSR signal>1.3 V; darker colors and bold line) for four different trial types. B. The release RTs across the subjects for the No Stop, Error Stop, and partial Error Stop trials shown in the plot for high and low starting forces indicate that the higher starting force leads to higher release RTs. C. The plot shows the effect of starting force on the FSR signal release velocity after the Go signal. Higher starting force results in faster decay in the applied force, however, in both initial conditions, the partial Error Stop trials were characterized by the lowest magnitude of release velocities. D. The figure shows the distributions of sensor-estimated SSRTs estimated independently for the high and the low starting force, which don’t exhibit a significant difference across the two conditions.

Indeed, the higher force on the mouse button was associated with longer RRTs across all the trial types, and the No-Stop RRTs were longer than the RRTs observed in Error Stop and partial Error Stop trials for both the force levels as indicated in Figure 8B. However, we did not detect a significant interaction between the two factors (F(2,18)=0.14; p=0.87). Additionally, we found that the force condition affected the velocity of the force decay after the Go signal. In both force conditions, the partial Error Stop trials exhibited the lesser velocity of FSR signal release compared to No-Stop and Error Stop trials; and the overall rate of decay in low force trials was of lesser magnitude (Fig. 7C). Finally, we tested the influence of the level of starting force on inhibition, and found no significant differences between the sensor-estimated SSRTs obtained from partial Error Stop trials across different force conditions (Fig. 7D; ttest t(9)=2.26; p= 0.49), thus confirming the observations of the previous experiment.

**Figure 8.**
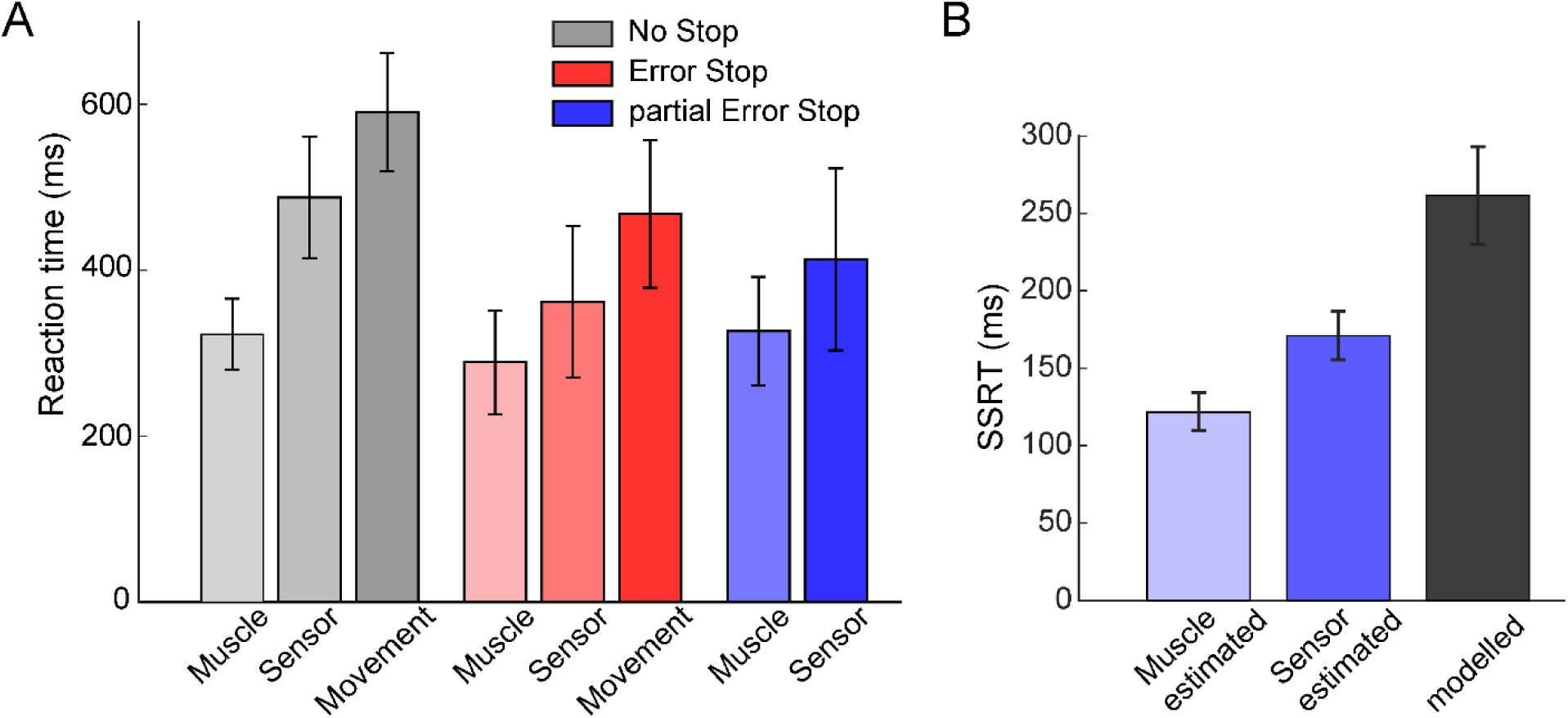
Comparison of motor behavior estimates from FSR signal and muscle activity. A. The figure shows a comparison between the reaction times detected by three different modalities, i.e., the muscle activity, the FSR signal, and the overt movement detected by the Button release. The height of the bars represents the means and the vertical bars represent the STD of the RTs and SSRTs across subjects (n=6). B. The plot compares the SSRTs obtained by the partial Error Stop trials detected from the muscle activity and the Force signal to the model-estimated SSRTs.

### Time course of the cascade of events occurring during motor execution and cancellation: Relationship between the effector force and EMG

In our previous analysis, we observed that the FSR sensor estimates of SSRT anticipated the behavior-based estimate of SSRT by about 88 ms. Similarly, in both the No-Stop and Error Stop trials, we observed that the RRTs detected by the FSR occurred about 120 ms before the overt displacement of the button following the Go signal. Previous studies have shown that signatures of action execution and cancellation can be detected in physiological signals such as muscle activity, well before evident limb movement (9, 16). In line with those investigations, we demonstrated a reflection of action control in the force modulation here.

We investigated how force modulation can be inserted into the cascade of events that lead from muscle activation to movement initiation. To this end, we recorded agonist EMG activity during finger displacement from six participants. Figure 8 illustrates the relationship between muscle activity and the dynamics of force release when the movement was either completely or partially executed. As expected, the muscle RTs preceded the sensor-based RRTs leading to movement execution, observed as the button release with even longer latencies; or the inhibition in the partial errors accompanied by no overt response (Fig. 8A). Similar patterns were also observed in the SSRTs obtained from the muscle activity and the FSR sensor signal dynamics (Fig. 8B), i.e., muscle-estimated SSRT (122±12 ms) preceded the sensor-estimated SSRT (171±14 ms) by 49 ms, while both of these estimates preceded the SSRTm (262±29 ms).

Despite this anticipated modulation in EMG compared to FSR, we found that we could detect the incoming response by employing the FSR signal in more trials compared to the EMG activity (978±22 vs 659±130). The same tendency was also observed when considering the partial Error Stop: FSR signal (42±15), EMG signal (18±4). Thus, even though the FSR detected the changes associated with the behavioral outcome later than the EMG signal, the amount of information obtained through the analysis of the force signal asserts it a better measure to predict the behavioral outcome during movement initiation and inhibition.

## DISCUSSION

We used a force sensitive resistor to explore the impact of initial finger force unconsciously applied on a mouse button on motor control during a Stop Signal Task (SST). Using the analog signal from the Force Sensitive Resistor (FSR) as a continuous measure of the applied force, we found that variations in force dynamics before movement initiation could influence overt movement occurrence. More specifically, the force level before the Go signal influenced both the latency of movement initiation and the possibility to cancel it, if needed. Two controlled force levels imposed in a separated experiment confirmed these observations.

Changes in force indicated movement inhibition in partial Error Stop trials, allowing estimation of the Stop Signal Reaction Time (SSRT) at single trial level. However, SSRT estimates did not vary with the level of applied force. Electromyography (EMG) activity preceded force release, but the force release was more reliable than EMG in detecting movement onset in partial Error trials, suggesting a stronger limitation of the surface EMG in monitoring single muscle activity for movement initiation or inhibition.

### Effector’s initial state affects overt behavior

In most studies that investigate motor control by employing the Stop Signal Task, the initial state has been considered more from the perspective of cognitive and strategic factors as proactive control (17–20), while here we introduced a biomechanical variable to be evaluated. It is well established that movement generation is typically associated with the coordination of different muscles and with muscle synergies (6, 7, 21) often contributing anticipatory postural adjustments when movements influence the body postural state (12, 14). Therefore, activation changes associated with the incoming action can potentially be detected at the level of the acting effectors rather than with the more relevant EMG agonist activation, as typically monitored. This is also true for focal (one-finger) movements where the coordinated activity of several hand muscles is observed (22). For example, the request to generate force with one finger can be accompanied by the generation of force in other fingers in a multi-finger task (23). All these data suggest that the overt “movement” observed can be preceded by a distributed and coordinated pattern of contractions that involves more muscles than the ones typically recorded, which in turn could be affected by biomechanical constraints, such as multi-joint tendons (24). This aspect is crucial as it requires a measurement that captures the coordinated activity of such muscles, especially considering that biomechanical postural variables potentially influence the Go and Stop processes. Here, we explored the force applied to the button as an output of this coordinated activity and found that higher force correlated with longer reaction times, as well as the occurrence of partial errors or correct stop trials.

However, the level of force did not produce any changes in SSRT in the experiments 1 and 2. Other studies reporting force measurements during an SST have found that changes in force during the movement execution (squeezing or keypress) were linked to changes in probability to respond or the inability to inhibit during stop trials (11, 25). However, no clear correlation is presented between the applied or baseline force and the stopping latencies. We attribute these findings to the differing nature of experimental designs. In a similar fashion, our results also revealed a higher probability to inhibit, if the trial was started with a higher baseline force before the Go signal.

In our study, the required amount of force was applied before the Go signal, rather than after it. Moreover, we did not request the subject to monitor the applied force. Under this condition, we speculate that the lack of modulation in the sensor estimated SSRT in our study may be due to the request in Stop trials to maintain the current state. Consequently, the force level may not have significantly impacted this parameter of inhibition. On the contrary, No-Stop trials required a state change, such as movement, which could have been more challenging under a higher force applied on the button. This condition could have impacted the probability to inhibit a movement without the need to change the SSRT, given the slower run of the Go process. This process is also reflected by the higher baseline force observed in the correct and partial Error Stop trials compared to the No-Stop trials, and furthermore a higher baseline force corresponding to more successful inhibition. To explore these differences in modulation, further studies requiring continuous movement interruption could be employed (26, 27).

We employed a method to detect changes in the background force, which can also identify signs of inhibition at the single trial level, as observed in partial Error Stop trials. Through analysis of the FSR signal, we estimated SSRT at the single trial level. Our analysis revealed that this estimate is shorter than SSRT obtained through behavioral analysis, as expected from signals preceding the behavioral response studies, and longer than those obtained through EMG activity, as evidenced in Figure 8. This highlights a detectable intermediate step between the onset of muscle activation and movement onset. Additionally, these values correlated with the behavioral estimate, as observed in recent studies involving finger (28) or arm movements version of SST (8, 9, 29).

Even observing a sequence of events consistent with previous studies here we detected some differences in the timing of their occurrence. For example, our muscle estimate of motor inhibition time occurred earlier than the one reported in key press versions of the task (123 vs 160 ms) and the key release time in Go trials is longer (about 570 ms vs about 486 ms) than in previous works testing motor inhibition through a stop signal or countermanding task (9, 16).

These differences can be accounted for by variations in the way the effector is required to move, in the recruited muscles, in the algorithms implemented to detect the changes, and in the mechanics of the devices used (30, 31). In our task the participants were required to release a button that they kept pressed until the Go signal, a condition entailing to contrast a higher amount of inertia than pressing a keyboard. This difference has likely prolonged the time of the button release. The earlier estimate of muscle SSRT it is possibly due to this experimental condition, that could have facilitated movement inhibition by making more difficult movement generation, and to the specifics of the algorithm used for the estimation. Indeed, we used a velocity-based algorithm implemented to capture the peak of velocity of the signal. Other works estimated the SSRT on the basis of the peak of the EMG signal. By applying this method, we obtained a further estimate of SSRT of 172 ± 24 ms, that is delayed by 52 ± 8 ms compared to the velocity estimate reported in our results, but is closer to the value of approx. 158 ms in previous works recently reviewed (16).

The possibility to obtain the SSRT at the single trial level is particularly important as it allows for advancements in understanding the intertrial variability of the inhibition process empirically, and it could also enable reliable SSRT estimation with fewer trials. Our findings suggest that a force sensor enhances the detection of this variable compared to monitoring single muscle activity. This approach appears to capture the muscle activity output, overcoming numerous technical challenges associated with signal acquisition, preprocessing, and analysis, as noted in previous EMG studies (8). This aspect is particularly useful when dealing with specific populations of subjects as patients or children. A further advantage of the signal we recorded is that it requires only the displacement of a force sensor on the mouse button, enabling a comfortable and ‘ecological’ interaction with the computer device. Therefore, we believe that its utilization could offer a valuable approach for obtaining the SSRT at the single-trial level compared to methods that necessitate rarer and less frequent movements (i.e., middle-lateral finger movements; (9, 32, 33) in order to effectively isolate the activity of the muscles involved.

### Evidence of a biomechanical Point-of-No-Return

The current findings align with previous observations indicating that the initiation of movement is preceded by changes in other physiological and biomechanical parameters that anticipate its overt manifestation (5, 9, 14). Different studies have shown that modulation in these parameters do not automatically lead to an overt movement. For instance, an increase in EMG response in agonist muscles during a proportion of correctly stopped trials suggests that partially initiated movements, in terms of EMG activity, could be halted before their overt manifestation. Overt manifestation occurs only if EMG activity surpasses a certain level. This implies that within the nervous motor somatic system, a certain level of activity must surpass the so-called ‘point of no return,’ beyond which the movement will be generated.

This activity is considered to be above the point of no return because if it is surpassed, it impedes the inhibition of movement despite the presentation of a Stop signal. In our research, we found a corresponding point of no return in the level of force applied to the sensors. Specifically, movements were generated when the level of force dropped below a certain threshold, as observed in No-Stop and Error Stop trials, but not in partial Error Stop trials. The drop in force could be considered as a signal reflecting the biomechanical state changes occurring in the hand, which are related to an increase in the activity of agonist muscles and a decrease in antagonist activity to generate the movement.

Thus, besides perceptual or cognitive variables (34–36), also the biomechanical factors can delay the run of the go process, a condition that increases the likelihood of a Stop signal successfully halting the movement (see 37 for review). In the present study, this condition was attributed to the higher force on the button, which decreased the velocity of force release and delayed the reaction time.

In a framework where initiating motor action begins with the neuronal processing of internal and external variables and proceeds with the recruitment of a set of muscle (9), our research underscores the significant influence of the initial biomechanical state, reflected in the amount of background force of the effector, which putatively influences the degree of muscle activation. Specifically, our findings reveal that greater force exerted on the mouse button due to initial state could delay the onset of force release and subsequent force release velocity. This delay increases the probability of interrupting a movement already in progress when a Stop signal occurs. Importantly, our study did not manipulate specific perceptual or cognitive variables, thereby excluding the possibility of a stage preceding muscle activity and force release that could impact the progression of the go process. Concurrently with the recent EMG findings, we have shown that the movement can be halted or cancelled until the very start of its execution, and, furthermore, that it is possible to investigate the inhibitory processes during a Stop-signal paradigm at a finer temporal scale when proper physiological or biomechanical measures are employed.

### Implications to motor inhibition modelling and physiological considerations

Combining the present results with previous studies, our data support the hypothesis that inhibition can occur at any time before the overt manifestation of a movement (5, 11, 25). In our case, this corresponds to the generation of sufficient muscle contraction to release the finger from the starting position, allowing the overt movement detection through mouse button release. As previously noted, this evidence challenges the assumption that the race between the Go and Stop processes is a winner-take-all mechanism, where the process that reaches the threshold first shuts down the other(11).

The observation that, after the stop signal is delivered, there is an attenuation of muscle or force-related activity associated with the Go process suggests that the Stop process is not limited to blocking muscle activity inputs. Rather, it continues to influence muscle activation even after the movement has been initiated. A key issue here is determining where the threshold should be placed in relation to goal-oriented movements. For example, if our task had required full finger extension, inhibition could have occurred before the completion of the movement, aligning with the predictions of the race model, as observed in reaching movements toward peripheral targets (8).

Our findings are consistent with the hypothesized interaction between the Stop and Go processes (11, 38), that as has also been recently observed in the inhibition of step initiation. In this case, inhibitory effects were reflected in the anticipatory postural adjustments, not only in partially initiated movements but also in correctly and erroneously stopped trials (14). These real-world situations are generally not fully captured by simplified experimentally controlled contexts and could not be accounted for by models of motor inhibition.

By broadening the perspective on inhibitory control, it can be hypothesized that inhibition occurs at different levels of the motor system depending on task demands and the timing of the inhibitory signal (5, 9, 39). Following this framework, inhibitory signals delivered early—such as at shorter SSDs—could inhibit motor programs at very early stages, during their maturation in the prefrontal (3) or premotor cortex (40, 41). In such cases, no muscle activation would be detected. On the other hand, if the inhibitory signal is delivered when the motor program is further along in its maturation, recruiting downstream motor areas, a reflection in muscle activity and force release should be observed. Accordingly, we observed this reflection for SSDs of intermediate and long durations. For the longest SSDs, we did not detect any peripheral evidence of motor inhibition, likely because we set the movement threshold at its initiation point. However, we found evidence of motor inhibition for intermediate SSDs, in trials with partial stop errors, suggesting an interaction between the Stop and Go processes. The estimated timing of inhibition based on force-sensing resistor (FSR) data aligns with the involvement of the basal ganglia hyperdirect inhibitory pathway on the motor cortex, as hypothesized for this rapid form of inhibition (9).

Furthermore, our findings introduce a new perspective on the beginning of the race process, highlighting that mechanical factors can influence the evolution of the Go process by delaying movement onset, similar to how cognitive and perceptual factors do (34, 42). This delay provides the Stop process with more time to successfully inhibit movement initiation.

Considering the latest context, how the maintenance of a starting (postural) position is managed by the nervous system requires further investigation. A possible field of investigation is examining the mechanisms of somatosensory gating, which may involve either spinal cord or cortical structures (43–46). This mechanism limits the access of non-relevant somatosensory information to the somatosensory cortex, allowing the motor system to prioritize sensory information relevant to the upcoming or ongoing movement. Supporting this hypothesis is evidence of the attenuation of somatosensory event potentials prior to movement initiation. The amplitude of this signal has been observed to correlate negatively with the degree of contraction of the agonist muscle (47, 48), or increase in response to the requirement to maintain a static position (49). In the present context, the lower degree of agonist contraction under high background force conditions, combined with the requirement to maintain a static position during stop trials, may have interfered with somatosensory gating and delayed movement reaction times (46). Measuring somatosensory event potentials during the execution of the proposed task could help clarify the extent to which interfering somatosensory information impacts motor inhibition and identify which neural structures are primarily involved in this process.

## DATA AVAILABILITY

The data that support the findings of this study are available from the corresponding author upon reasonable request.

## ACKNOWLEDGMENTS

We are grateful to Prof. Sabrina Fagioli for technical and theoretical support. We also thank Dr. Stefano Colangeli for contributing to data acquisition.

## AUTHOR CONTRIBUTIONS

EB and PP conceived and designed research, interpreted results of experiments, drafted manuscript, edited and revised manuscript, approved final version of manuscript. SR performed experiments, analyzed data, interpreted results of experiments, prepared figures, drafted manuscript, edited and revised manuscript, approved final version of manuscript. SF interpreted results of experiments, edited and revised manuscript, approved final version of manuscript. IBM,FDB and GB, edited and revised manuscript, approved final version of manuscript.

